# Sustained visual signals in the primate cerebellar dentate nucleus drive associative learning

**DOI:** 10.1101/2025.04.13.648306

**Authors:** Yusuke Akiyama, Hiroshi Yamada, Masayuki Matsumoto, Jun Kunimatsu

**Affiliations:** Graduate School of Comprehensive Human Sciences, University of Tsukuba, Tsukuba, Ibaraki 305-8575, Japan; Division of Biomedical Science, Faculty of Medicine, University of Tsukuba, Tsukuba, Ibaraki 305-8577, Japan; Transborder Medical Research Center, University of Tsukuba, Tsukuba, Ibaraki 305-8577, Japan; Center for the Evolutionary Origins of Human Behavior, Kyoto University, Kanrin, Inuyama, Aichi 484-8506, Japan

## Abstract

A number of studies have suggested that the cerebellum has cognitive functions; however, the underlying neuronal mechanisms remain unclear. In this study, we demonstrated that sustained visual signals in the cerebellar dentate nucleus represent the visuomotor associative information. We recorded neural activity from the dentate nucleus when monkeys performed a learning task involving the association between visual objects and saccade directions. We found that sustained visual activity was greater in the region during learning than during memory retrieval. This enhancement disappeared under the uncertain reward condition, wherein the monkey did not perform the heuristic learning strategy. Furthermore, sustained visual signals changed the response to visual objects depending on the associated saccade direction. This direction selectivity was positively correlated with modulation during learning. These results suggest that sustained visual signals in the dentate nucleus reflect learning motivation and drive learning by increasing the strength of discrimination among visual objects.

## Introduction

Several studies have suggested that the cerebellum contributes to various non-motor functions. Brain activity in the cerebellum has been linked to motivation^1, 2^, attention^3, 4^, social behaviors^5, 6^, and various emotional behaviors^7, 8^. The cerebellum has been implicated in psychiatric disorders such as autism spectrum disorder and attention-deficit/hyperactivity disorder^9, 10, 11, 12, 13^, suggesting that these disorders may be at least partially caused by impairments to its function. Therefore, understanding the neuronal processing of these non-motor cerebellar functions is critical. However, because the traditional focus in cerebellum research has been on its role in motor control and execution^14, 15^, it remains unclear which signals related to non-motor cognitive functions are processed in the cerebellum.

It was recently reported that the cerebellum, cerebral cortex, and basal ganglia are interconnected and involved in a number of common functions^16^. Although reward-based learning has mainly been investigated in the basal ganglia circuit, reports have indicated that the input to the cerebellar cortex contains reward-related information as well^17, 18, 19, 20, 21^. Clinical studies have reported that non-motor associative learning is impaired in patients with cerebellar degenerative diseases^22, 23^. Ipata et al. reported that neural activity in the mediolateral cerebellar cortex (crus I and II) changes in response to rewards during the process of visuomotor associative learning^24^. This activity has been shown to be connected to the medial prefrontal cortex. However, previous studies on visuomotor associations in the lateral prefrontal cortex and striatum have reported that sustained activity during the delay period encodes information related to learning^25, 26^. The cerebellar dentate nucleus, which has strong connections to the lateral prefrontal cortex, may also be involved in associative learning.

Kunimatsu et al. found that sustained signals in the dentate nucleus during motor preparation were related to motor strategies^27^. Similar preparatory activities also regulate higher-order control of movement, such as its timing^28^, self-generated movement-related behavior^29^, and motor planning^30^. Previous neurophysiological reports have shown that sustained activity, rather than phasic, is related to cerebellar function, indicating that sustained signals regulate the processing of cognitive information through the cerebral cortex.

These observations prompted us to explore sustained signals in the dentate nuclei of monkeys during associative learning. In this study, we recorded neuronal activity from single neurons in the dentate nuclei of monkeys and observed sustained visual signals while they were performing learning tasks regarding the associations between visual objects and motor directions. These sustained visual signals encoded the learned visual objects, and were enhanced during learning but not during memory retrieval. This enhancement disappeared under the uncertain reward condition, wherein the monkeys did not use the appropriate learning strategy—indicating that the enhancement of sustained visual responses occurred specifically when the monkeys were willing to learn the association. Thus, the enhancement of sustained visual signals may reflect the motivation or intention to learn, and facilitate learning by increasing the degree of discrimination between different visual objects.

## Results

### Behavioral strategy during visuomotor associative learning

Two monkeys performed a visuomotor associative learning task that required them to learn the association between two visual objects, involving both leftward and rightward saccades (Figure 1A). W each monkey looked at the fixation point, one of two fractal objects was presented in the center of the monitor. The saccade targets were then presented at left and right positions on the monitor. The monkeys received a liquid reward if they chose the correct target associated with the fractal object. One of the two fractal objects was randomly selected for each trial. The task was performed under two conditions (Figure 1B): the learning condition and the over-trained one. Under the learning condition, we used novel sets of fractals for each session, comprising a total of > 80 trials. The correct rate gradually increased with the number of trials (Figure 1C, red lines). By contrast, in the over-trained condition, the monkeys had previously learned about the association for > 10 sessions, so they maintained a high correct rate from the beginning of each session (Figure 1C, blue lines).

**Figure 1.**
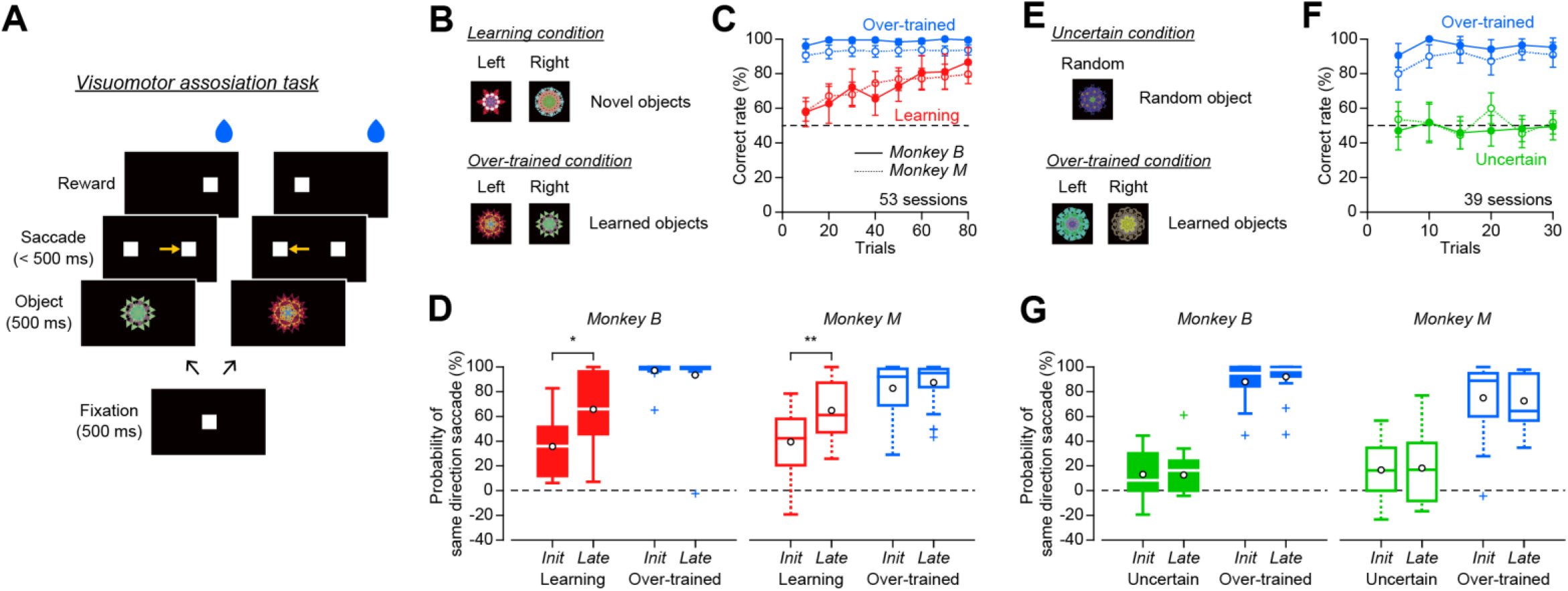
Visuomotor associative learning task and learning strategy. (A) Sequence of events during the visuomotor associative learning task. Following the white fixation point, one of two fractal objects was presented in the center of the screen. Two white target spots appeared to the left and right at the time of the object’s offset. The monkey then chose one of the targets through its eye movements. The correct target was associated with the fractal object. (B) Under the learning condition, a set of novel objects was used. Under the over-trained condition, a set of learned objects that the monkey has experienced > 10 sessions was used. (C) Correct rates in the learning and over-trained conditions. The mean correct rate and 95% confidence interval for every 10 trials were plotted. The solid and dashed lines indicate the data from monkeys B and M, respectively. (D) The difference in probability of a saccade in the same direction when the same object was presented in the center after correct and erroneous responses. Trials in the first and second half of each session were conducted as initial (Init) and late phases (Late), respectively (Wilcoxon rank-sum test, \*\**p* < 0.01). (E) Under the uncertain condition, a set of learned objects were used along with one random object. (F) Correct rates for the learned objects and random one. (G) The difference in probability of a saccade in the same direction when the same object was presented in the center after correct and erroneous responses.

To examine the monkeys’ behavioral strategies during the learning condition, we analyzed their behaviors following both correct and erroneous trials for the presentation of the same object. If they exhibited heuristic behavior based on the reinforcement learning theory, we predicted that they would make saccades in the same direction to the same object after a correct trial, and in the opposite direction after an erroneous one^31^. We calculated the difference in the probabilities of the monkeys making saccades in the same direction as the one it was originally presented in when the same object was presented, after both correct and erroneous trials, and compared it between the initial and late phases under each condition (Figure 1D). In the learning condition, the probability differed significantly between the initial and late phases in both monkeys B (Figure 1D left, Wilcoxon rank-sum test *p* = 0.0038) and M (Figure 1D right, *p* = 3.1 × 10^-4^), but no difference was observed in the over-trained condition (monkey B *p* = 0.62; monkey M *p* = 0.48), indicating that the monkeys’ behaviors adapted to a win-stay lose-shift strategy as the learning progressed.

During the initial learning phase, the probability of reward was uncertain. This reality may have influenced activity in the dentate nucleus activity. Therefore, as a control for the learning condition, we also used the uncertain condition—wherein a fractal object was not associated with any saccade direction (Figure 1E, random object), and the correct saccade direction was changed randomly. The monkeys experienced this condition over 10 sessions, similarly to what occurred in the over-trained condition. The monkeys performed this condition by mixing it with a set of learned objects wherein the correct saccade direction was fixed. The correct rate was consistently high for the learned objects during this test, whereas it was at a random chance level rate for the random object, since the monkeys could not predict the correct saccade direction (Figure 1F). Moreover, the behavioral strategy toward the random object was nonoptimal across all phases (Figure 1G, monkey B *p* = 0.88; monkey M *p* = 0.95), indicating that the monkeys were not motivated to learn the association for it.

### Two types of visual signals in the dentate nucleus

We recorded the activities of single neurons in the posterior region of the cerebellar dentate nucleus while the monkeys performed the visuomotor associative learning task. We focused on their response to a fractal object in this study because the ability to integrate temporally separated stimuli and response information is particularly important in associative learning, and neural activity in response to cue stimuli is thought to play a role in this process within the prefrontal cortex^32^. A previous study reported that activity in the cerebral cortex during the delay period following visual object presentation contributed to associative learning^25, 26^. Principal component analysis revealed two components of the visual response: the early phasic and the late sustained response (Figure S1). Based on this analysis, we defined the phasic-response neurons as those that exhibited activity during the early visual period (between 1–150 ms immediately after object presentation) and the sustained-response neurons as those that exhibited activity during the late visual period (between 201–400 ms after object presentation). We found 56 phasic-response neurons (27 and 29 neurons in monkeys B and M, respectively) and 181 sustained-response neurons (101 and 80 neurons in monkeys B and M, respectively) in the ventral and dorsal regions of the dentate nucleus (Figure 2). Because we did not find a clear difference in neuronal activity between the ventral and dorsal groups, the data were combined for our subsequent analyses. A total of 32 neurons were classified as both phasic- and sustained-response neurons, but were plotted as sustained-response neurons in Figure 2.

**Figure 2.**
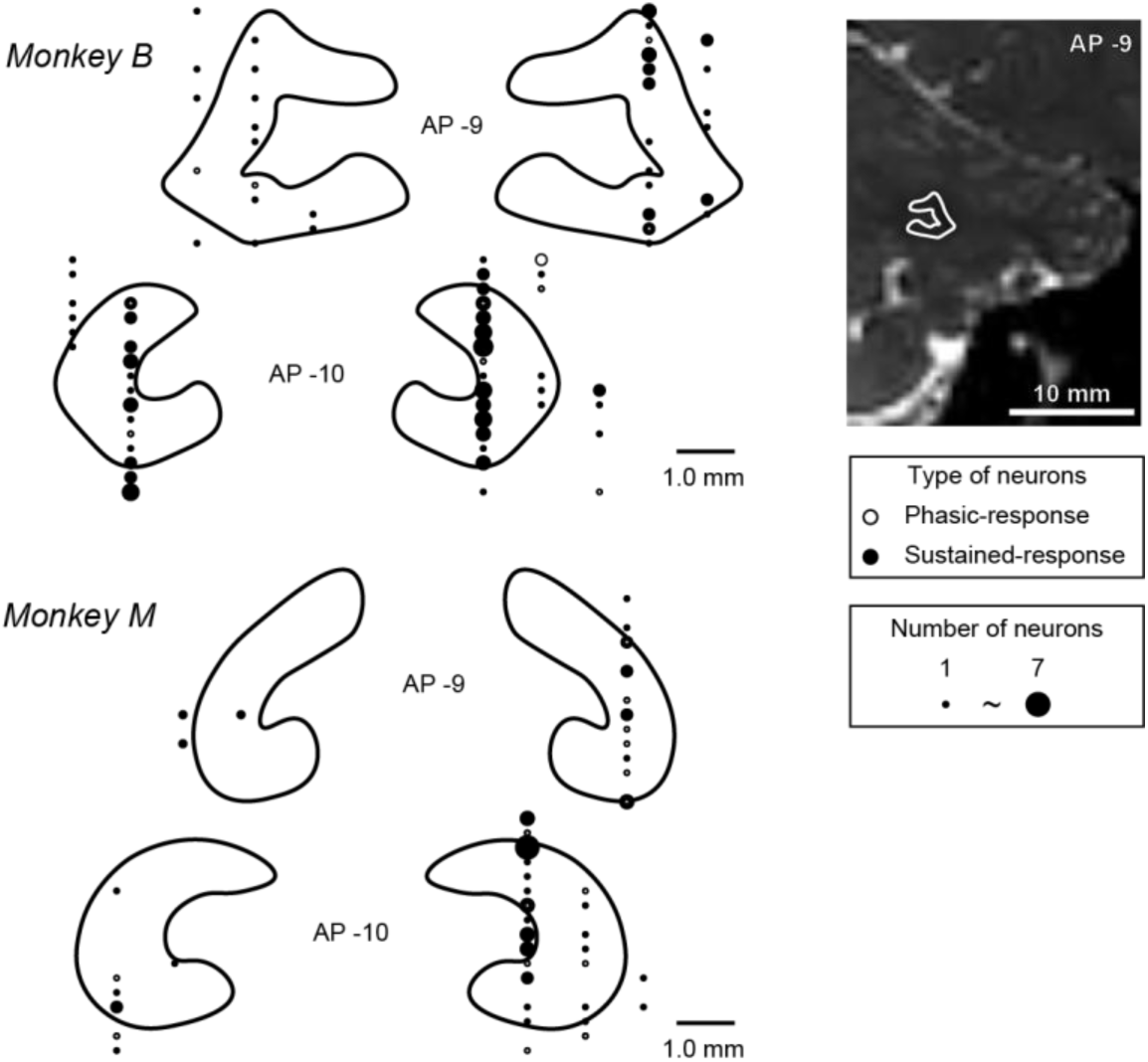
Locations of recording sites. Coronal sections showing the locations of visual neurons (filled dots, sustained-response; unfilled dots, phasic-response). These sections were taken 9–10 mm posterior to the anterior-posterior (AP) and reconstructed via MR imaging (T2-weighted image, top-right panel). Neurons with both sustained and phasic response characteristics were grouped together as sustained-response neurons. The size of the circle represents the number of neurons. Some neurons from the AP-9 and AP-10 regions were also plotted alongside those from the AP-8 and AP-11 ones.

To examine the changes for each response type that take place during visuomotor associative learning, we compared the neuronal responses between the learning and over-trained conditions. Figures 3A and 3B show the firing rates for representative phasic- and sustained-response neurons. The phasic-response neuron was equally active for both novel and learned objects (Figure 3A; unpaired-samples Student’s t-test *p* = 0.54), whereas the sustained-response neurons were active only for novel objects (Figure 3B, *p* = 1.1 × 10^-14^). Thirty of the 56 phasic-response neurons showed significant differences between the learning and over-trained conditions during the early visual period (unpaired-samples Student’s t-test *p* < 0.05; filled circles in Figure 3C), and there was no significant difference in the population (paired-samples Student’s t-test *p* = 0.14). The time courses of the population activity for the 56 phasic-response neurons also showed no differences between the learning and over-trained conditions (Figure 3E). By contrast, 97 of the 181 sustained-response neurons showed significant differences during the late visual period (*p* < 0.05; filled circles in Figure 3D). The activity in this population was significantly greater during the learning condition vs the over-trained one (Figure 3D, *p* = 7.1 × 10^-7^). The time courses of the population activity for the 181 sustained-response neurons confirmed that the activity during the learning condition was greater than that during the over-trained condition (Figure 3F).

**Figure 3.**
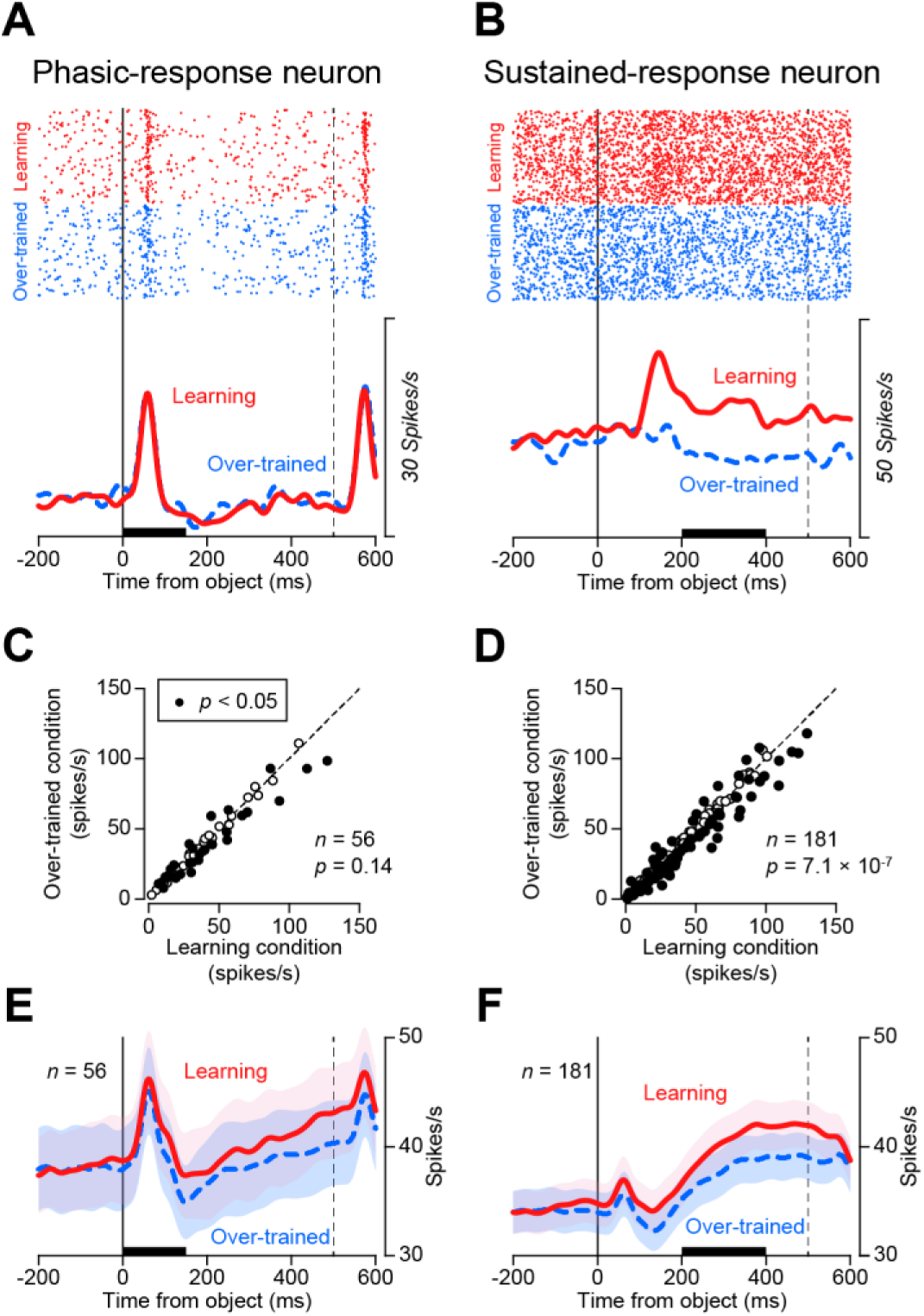
The phasic and sustained visual responses under learning and over-trained conditions. (A) An example of a phasic-response neuron. Data are aligned on the object onset (vertical line). Red solid and blue dash lines indicate the activity levels during the learning and over-trained conditions, respectively. Dashed vertical line indicates the onset of target spots presentation. (B) An example of a sustained-response neuron. (C) Each data point compares the mean firing rate during the learning condition with that during the over-trained one, for each phasic-response neuron. Mean firing rates were measured from the early visual period (black bar in A, 1–150 ms after object presentation). Filled symbols indicate the data, showing a statistically significant difference between the conditions. (Wilcoxon rank-sum test, *p* < 0.05). (D) Each data point compares the mean firing rates during the learning condition with those during the over-trained one, for each sustained-response neuron. Mean firing rates were measured from the late visual period (black bar in B, 201–400 ms after object presentation). One outlier point has been omitted (165.9, 162.3), but is included in the histogram. (E,F) Time courses of the population activity levels during the learning and over-trained conditions for the phasic- (E) and sustained-response neurons (F). The traces indicate the mean and standard error of spike density for individual neurons.

### Sustained visual signals during the uncertain condition

The sustained visual responses were enhanced under the learning condition. What does the sustained signal represent? Recent studies have reported that neurons in the septum and striatum show strong responses to uncertain reward cues, but not to the ones that indicate certain outcomes^33, 34^. Sustained visual signals in the dentate nucleus may also encode the context of uncertain reward. To examine this, we investigated the activity levels of the sustained-response neurons we identified under the uncertain condition (Figure 1E), which involved one random object and two learned objects. The behavioral data under this condition revealed that the monkeys did not attempt to learn the association for the random object (Figures 1F and 1G). This suggests the monkeys knew that the random object was not associated with the specific saccade direction, likely through their previous experiences.

We examined the activity levels of the 152 sustained-response neurons under the uncertain condition, with a representative one shown in Figure 4A (the same neuron shown in Figure 3B). This neuron showed significantly greater activity during the learning condition vs the over-trained one (Figure 3B); however, it exhibited no significant differences in activity between the random and over-trained objects (which were associated with the ipsilateral or contralateral saccade direction, respectively; Figure 4A, one-way analysis of variance [ANOVA]). Among the 152 sustained-response neurons, 85 showed a significant difference between the three fractal objects (*p* < 0.05). We compared the responses to the three object types among the 152 sustained-response neurons quantitatively, but found no significant differences (Figure 4B, one-way ANOVA). This result was confirmed by a time course of the activity of the overall sustained-response neuron population (Figure 4C). These results suggest that the sustained visual response may not reflect the probability of an uncertain reward, but may be specifically enhanced when monkeys are motivated to learn through visuomotor association.

**Figure 4.**
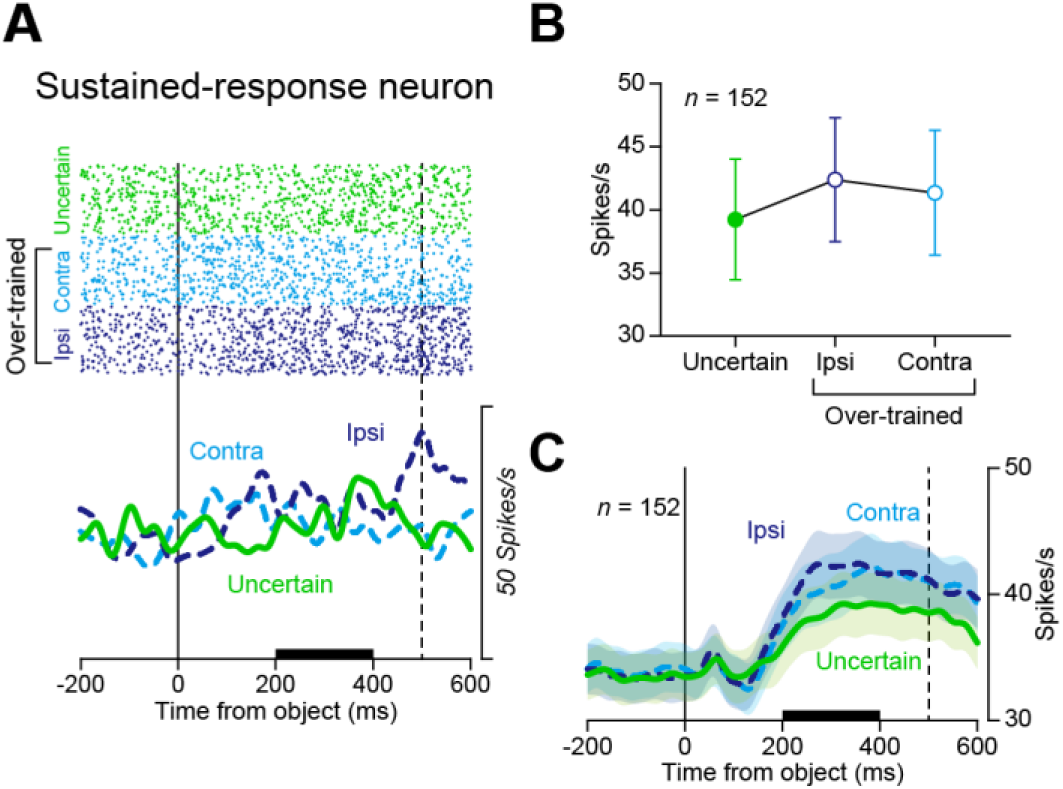
Activity of sustained-response neurons during the uncertain condition. (A) Example of the response of a representative sustained-response neuron (the same one shown in Figure 3B) to a random object (green solid line), an ipsilateral saccade object (dark-blue dashed line), and a contralateral saccade object (light-blue dashed line). (B) The differences in responses among three objects (one-way ANOVA, *p* = 0.66). Mean firing rates and 95% confidence intervals were measured from the late visual period (black bar in A). (C) Time courses of the activity of the overall sustained-response neuron population during the uncertain condition. Traces indicate the mean and standard error of spike density for individual neurons.

### Sustained visual signals encode visual objects based on the associated saccade direction

If cerebellar dentate nucleus neurons are involved in associative learning, they may do so by encoding task-relevant information. In our visuomotor associative learning task, the monkeys needed to determine whether to make saccades to the left or right direction when the object was presented. We compared the firing rates of the phasic- and sustained-response neurons response to the objects associated with saccades in both the ipsilateral (ipsilateral saccade objects) and contralateral (contralateral saccade objects) directions. Among the 56 phasic-response neurons, only two showed significant differences in their response to each object during the early visual period (unpaired-samples Student’s t-test *p* < 0.05). By contrast, 71 of the 181 sustained-response neurons showed significant differences in their responses to each object during the late visual period. We observed two types of responses: ipsilateral- and contralateral-object preferred response. Figures 5A and 5B show the activity levels of the sustained-response neurons, highlighting the ipsilateral- and contralateral-object preferred responses during the correct trials in the learning condition, respectively. In those trials (red), the preferred response was significantly stronger for the ipsilateral saccade object (solid line) vs the contralateral one (dashed line; Figure 5A, paired-samples Student’s t-test *p* = 1.7 × 10^-11^). Notably, no significant difference was observed in the number of erroneous trials (black, *p* = 0.35). The neurons with preferred responses to the contralateral saccade object showed the same results (correct, *p* = 1.5 × 10^-8^; error, *p* = 0.55). These results suggest that the sustained visual signal encoded the correct saccade direction based on visual stimuli, but was not related to motor preparation in a specific direction.

**Figure 5.**
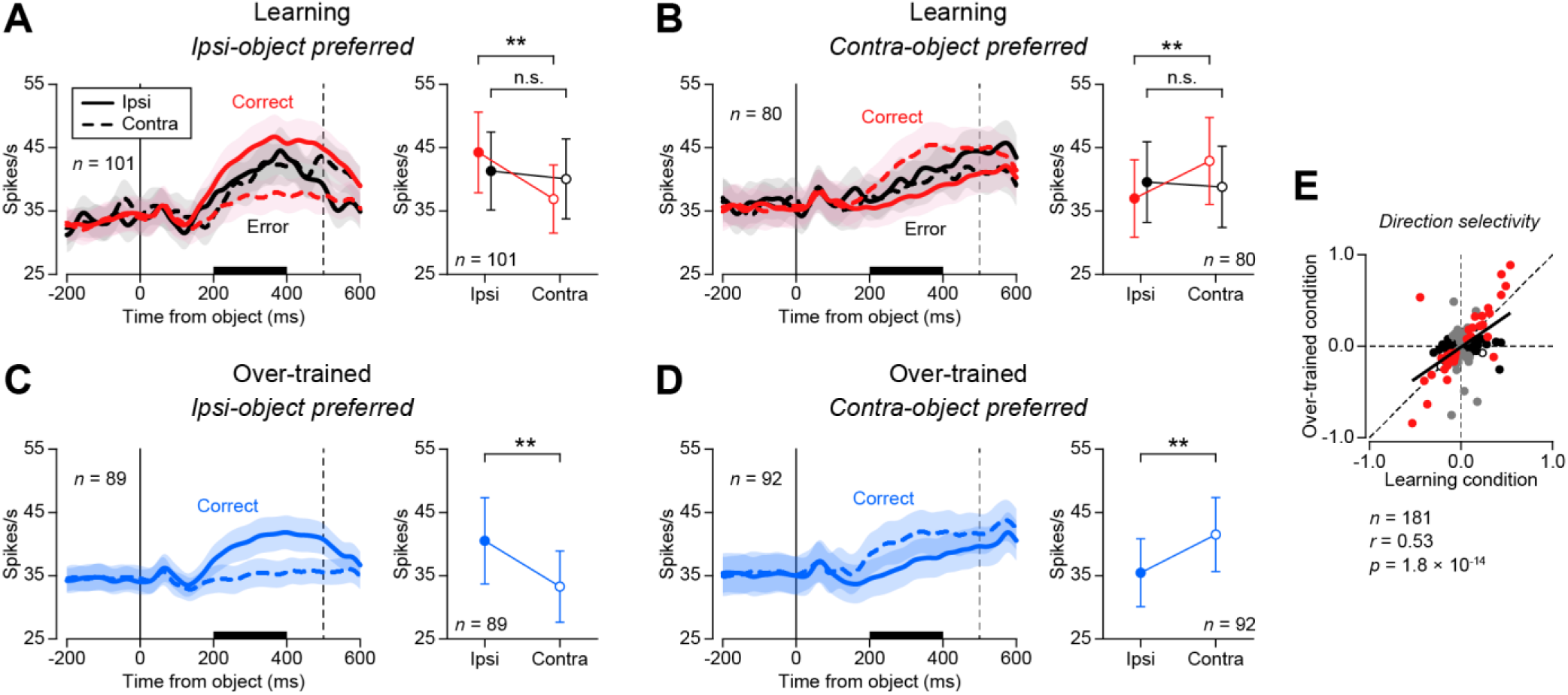
Direction selectivity of the sustained visual response. (A,B) Time courses of the population-level activity of the ipsilateral- (A) and contralateral-object preferred neurons (B) during the learning condition (left panel). Red and black lines indicate the responses to the ipsilateral (solid) and contralateral saccade objects (dashed) in the correct (red) and erroneous trials (black). Mean firing rates and 95% confidence intervals during the late visual periods following the presentation of ipsilateral and contralateral saccade objects (right panel, paired-samples Student’s t-test \**p* < 0.05; \*\**p* < 0.01). (C,D) Time courses of the population-level activities of the ipsilateral- (C) and contralateral-object preferred neurons (D) during the over-trained condition. (E) Correlation of direction selectivity within the sustained visual response between the learning and over-trained conditions. The color of each data point indicates neurons that exhibited different responses to ipsilateral and contralateral saccade objects (red: both the learning and over-trained conditions; black: only during the learning condition; gray: only during the over-trained condition; unfilled: no difference). The black line represents a linear regression line-of-best-fit.

We also examined the ipsilateral- and contralateral-object preferred responses in correct trials during the over-trained condition. Notably, the number of erroneous trials during the over-trained condition was small and insufficient for this analysis. In the over-trained condition, significant differences in sustained visual activity were observed between the ipsilateral- and contralateral object-preferred neurons (Figures 5C and D; 89 ipsilateral-object preferred neurons, *p* = 3.6 × 10^-8^; 92 contralateral-object preferred neurons, *p* = 7.7 × 10^-11^), similarly to the learning condition. To examine the relationship of the direction selectivity between the learning and over-trained conditions, we plotted the direction index during the late visual period for the correct trials (Figure 5E, see Methods). We found a positive correlation between direction selectivity under the two conditions (*r* = 0.53, *p* = 1.8 × 10^-14^). These results indicate that the encoding of object selectivity during learning was maintained after it had finished.

### Relation between direction selectivity and the enhancement of sustained visual signals

We proceeded to examine the role of sustained visual signal enhancement during learning, as this enhancement may affect the strength of direction selectivity. To examine this, we calculated the learning-modulation index by sustained visual activity under both the learning and over-trained conditions (see Methods), and measured its correlation with direction selectivity. Among the 181 sustained-response neurons, 39 and 46 showed both enhancement of sustained responses (unpaired-samples Student’s t-test *p* < 0.05) and clear direction selectivity (unpaired-samples Student’s t-test *p* < 0.05) under the learning and over-trained conditions, respectively. In Figure 6, we plotted the learning-modulation indexes and direction selectivity of these neurons. They exhibited a positive correlation between them under both the learning (left panel, *r* = 0.59, *p* = 8.3 × 10^-5^) and over-trained conditions (right panel, *r* = 0.66, *p* = 5.2 × 10^-7^). These results suggest that direction selectivity is regulated by the enhancement of sustained visual responses during learning, and is maintained afterward.

**Figure 6.**
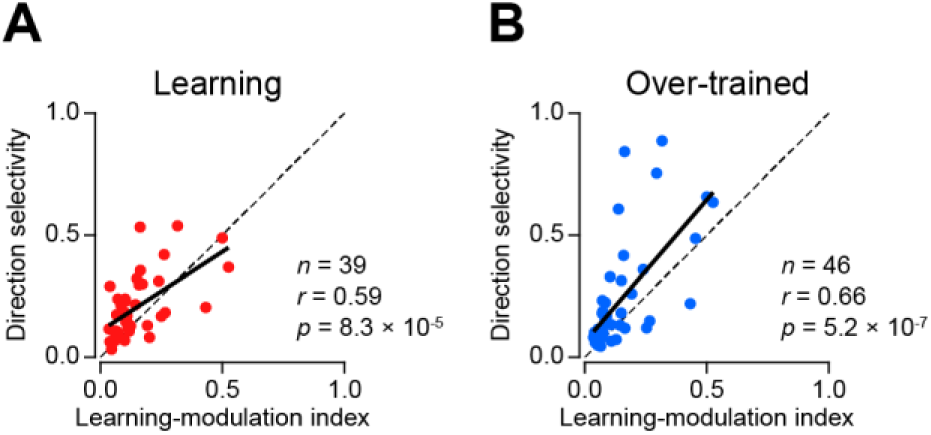
Correlations between learning-modulation index and direction selectivity in sustained-response neurons. (A) Correlation between the learning-modulation index and direction selectivity for each sustained-response neuron that showed significant learning-modulation activity and direction selectivity under the learning condition. Direction selectivity is shown as an absolute value, indicating selectivity for either the ipsilateral or contralateral direction. The black line represents a linear regression line-of-best-fit. (B) Correlation between the learning-modulation index and direction selectivity for each sustained-response neuron that showed significant learning modulation activity and direction selectivity under the over-trained condition.

## Discussion

In this study, we revealed that sustained visual signals in the dentate nuclei of monkeys represent information related to associative learning in two ways. First, the sustained visual signal represented the learning state (Figure 3) but not the uncertain situation (Figure 4). Second, the sustained response encoded the saccade goal direction, associated with visual objects during and after learning (Figure 5). Notably, this direction selectivity was stronger according to the strength of enhancement of sustained visual activity during learning (Figure 6). These results suggest that the enhancement of visual signals modulates learning by increasing direction selectivity.

Then, how does the neuronal circuit comprising the learning signal affect the direction selectivity? Figure 7 presents a diagram of a hypothetical neuronal network for visuomotor association learning. Our data suggested that the dentate nucleus neurons of our test monkeys were excited by novel visual objects during the learning condition (Figure 3). As we always used a novel object set during the learning condition, the increment of activity change may represent a response to the object’s novelty. However, novelty-related responses are typically phasic and diminish over several trials^35, 36^, making them unlikely to account for the observed activity in this experiment. However, the reward probability was low at the beginning of the learning phase, raising the possibility that this activity reflected the uncertainty of the reward, as reported in the septum and caudate nucleus. However, our results rejected this possibility (Figure 4). Rather, the fact that neuronal activity tended to be greater during the over-trained condition (without a significant difference) suggests that the monkeys were still attempting to learn during this phase. Thus, our results suggest that late visual signals represent the intention or motivation to learn about the associations between novel objects and movements. Previous studies have demonstrated that motivation is essential for learning^37, 38^, and imaging research has suggested that the cerebellum plays a role in learning-related motivation^39^. In rodents, cerebellar lesions have been reported to reduce learning motivation^1^. Based on these findings, we hypothesized that information indicating the learning state may originate from the cerebellar cortex (Figure 7).

**Figure 7.**
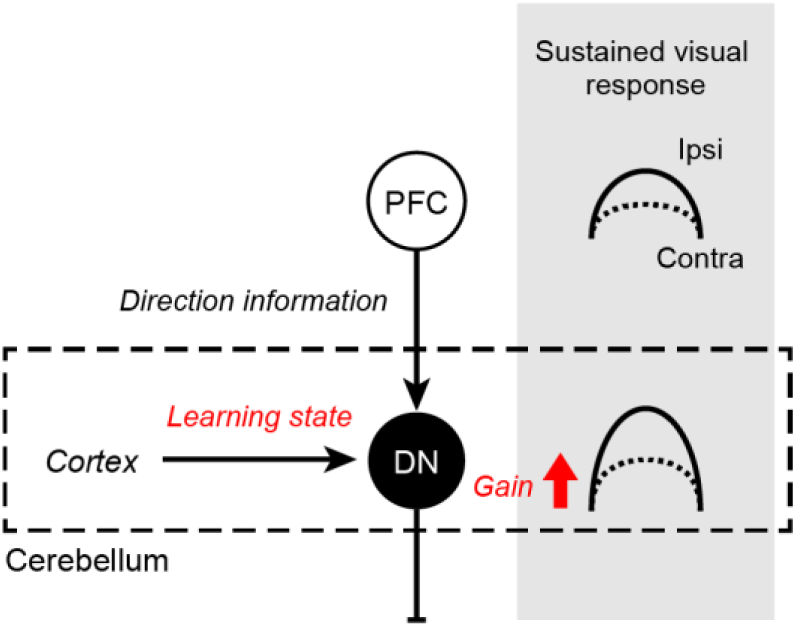
Hypothetical diagram of the neuronal mechanism underlying visuomotor associative learning in the cerebellum. Neurons in the dentate nucleus receive saccade direction information from the prefrontal cortex and signals that encode the learning state, from the cerebellar cortex. Sustained visual activity increases during learning and enhances the difference between responses to ipsilateral and contralateral saccade objects. DN: dentate nucleus; PFC: prefrontal cortex.

Although direction-selective signals during saccade preparation have been widely reported in various brain regions^40, 41^, our study found that such activity was not relevant to the dentate nucleus. Instead, our data reflected the direction of the goal associated with visual stimuli (Figure 5). This suggests that the dentate nucleus represents integrated information that combines visual objects and movement direction, which represents an essential process for associative learning. To the best of our knowledge, similar activity has not been previously reported in the literature regarding the cerebellum. However, direction-selective activity during visuomotor learning has been reported in the lateral prefrontal cortex (LPFC) and caudate nucleus^42^. The activity described by Histed et al. does not rule out a motor-related component^26^, but its time course closely resembles what we observed in this study—suggesting a functional connection between them. Indeed, a subset of the caudal dentate nucleus (the dorsal region, specifically) has been shown to project to the LPFC^43^. Moreover, recent studies have reported that the dentate nucleus receives direct cortical projections^44^. It is therefore possible that the posterior dentate nucleus has a reciprocal connection with the LPFC to aid in the integration of visual and movement-related information.

The increases in direction selectivity we observed during learning (Figure 6) suggest that the motivation/intention signal modulates visuomotor integration into the dentate nucleus. Although motivation changes behavior, the neural mechanisms underlying how motivation facilitates learning remain unclear. The difference between the contralateral and ipsilateral saccade object responses increased in this study, through the enhanced neuronal activity, which correlated with improvements in visual discriminability. A similar modulation of the visual response was reported in the visual cortex by Treue and Trujillo^45^. They found that attention increased neuronal responses to all stimulus directions, not just to stimuli moving in a preferred direction, which further improved discriminatory ability. These findings suggest common mechanisms by which internal states modulate the gain of the visual response. Notably, the discriminability of the test subjects regarding the visual objects was maintained in our study even after learning had already been established (Figures 5 and 6). This result suggests that sustained visual signals alter synaptic weights to maintain visual discrimination ability. To verify this hypothesis, the time-courses of neuronal activity levels in the frontal cortex and cerebellum must be explored in future studies. Furthermore, pathway-specific manipulation using pharmacological or optogenetic techniques would likely also significantly further the current understanding of these processes.

In behavioral tasks that require multiple processes, such as visuomotor learning, the brain often employs proactive control strategies. This is crucial for higher-order motor regulation, and several recent reports have described this type of sustained activity in the cerebellum^46^. Sustained activity has also been implicated in internal timing control^29, 47^ and motor planning^30^, as well as in behavioral strategy activities such as the speed-accuracy trade-off^27^. In this study, we found that a sustained visual response was active when the monkeys performed a suitable learning strategy (Figures 1 and 3). Sustained activity in the cerebellum may contribute to the proactive control of behavioral strategies at the cognitive and motor levels. Recently, Sendhilnathan et al. recorded reward-related signals in the cerebellar crus I and II^48^ during visuomotor association learning, and found that these signals were tonically active across trials. However, we did not observe such activity in the posterior dentate nuclei of our test subjects. This suggests that our study may have captured information from distinct neural pathways through crus I and II. Specifically, they demonstrate that crus I and II have strong connections with the medial PFC^49^, whereas the posterior dentate nucleus sends signals to the LPFC—suggesting that these signals are processed through different pathways. These visual and reward-related signals may be integrated within the frontal cortex to regulate visuomotor associative learning.

Thus, our study suggests that visuomotor association is achieved through the coordination of sustained activity between the PFC and dentate nucleus. To our knowledge, this represents the first proposed mechanism through which motivation or intention facilitates learning.

## Methods

### Animal preparation

Two Japanese monkeys (*Macaca fuscata*; monkey B male, 7.6 kg, 5 years old; monkey M, female, 6.6 kg, 5 years old) were used in this study. The care of the monkeys and all experimental procedures were approved in advance by the Animal Experimentation Committee of the University of Tsukuba (approval number: 20-266), and all experiments were performed in accordance with the protocol of the BioResource Project “Japanese Monkeys” of the MEXT, Japan. Prior to the study, we surgically placed a head-fixing device and recording chamber on each monkey’s skull, under general anesthesia (isoflurane, 0.5–2.0%). Behavioral task training and recording experiments were performed after the monkeys had fully recovered from surgery.

### Behavioral task

We controlled the behavioral task using MonkeyLogic^50^, which is the MATLAB (Mathworks) application. Each monkey sat on a primate chair with its head fixed in place, within a dark and soundproofed room. The visual stimuli used in the task were presented on a 24-inch monitor (ProLite E2483HS, iiyama) placed 42 cm in front of the monkey’s eyes.

We used a visuomotor associative learning task in which subjects were required to learn the association between two visual objects and leftward or rightward saccades (Figure 1). In each trial, a white square (0.5° × 0.5°) was presented in the center of the monitor, and the monkey was required to fixate on it for 500 ms. One of a pair of fractal objects (∼ 6.0° × 6.0°) was then presented. After fixating on the fractal object for 500 ms, two white targets (0.5° × 0.5°) were simultaneously presented in the left and right positions, at 13.5° from the center. The monkey was required to make a saccade to either the left or right target within 500 ms. If it fixated on the square in the correct direction, a high-pitched beep (2000 Hz) sounded and the monkey received a liquid reward. Otherwise (i.e., if the monkey made a saccade in an erroneous direction, did not make a saccade within the time window, or did not fixate on the target for the required duration), a low-pitched beep (200 Hz) sounded without a reward. A set of two different fractal objects was prepared for each session, and one object was presented randomly for each trial. One session consisted of 80–160 trials. The fractal objects used in these tasks were created using fractal geometry^51^.

The task was performed under three conditions: 1) In the learning condition (Figure 1B, top), we used a set of two novel objects for each session (novel object); 2) In the over-trained condition (Figure 1B, bottom), we used a set of two objects that the monkeys had already been trained on for > 10 sessions (learned object); and 3) In the uncertain condition (Figure 1E), we used an object in which the goal saccade direction was randomly changed in each trial (random object). Similarly to the over-trained condition, the uncertain condition was trained for > 10 sessions. A random object and a set of learned objects were included in each session.

### Recording procedure

Based on the stereotaxic atlas^52^, the recording chamber was positioned above the cerebellum of each subject. Magnetic resonance (MR) images (3T, Philips) were obtained along the direction of the recording chamber, which was visualized by gadolinium filling the grid holes. To record the activity of a single neuron, a stainless steel probe (S-probe, Plexon) was inserted through the guide tube. The insertion depth of the probe was controlled using a micromanipulator (MO-97, Narishige), and the recording position was controlled using a grid system fixed to the chamber. The probe was lowered into the cerebellar dentate nucleus (8–11 mm posterior to the interaural line and 6–9 mm lateral to the midline). The neuronal signals were amplified and filtered at a sampling rate of 40 kHz (OmniPlex, Plexon). The recorded neural activity data was separated into clusters using dedicated software (Offline Sorter, Plexon) to identify the activity of individual neurons.

### Data acquisition and analysis

Eye movements were monitored using an infrared eye-tracking system (EyeLink, SR Research) with sampling at 1 kHz. The timestamps of task events were concurrently sampled at 1 kHz and saved as files during the experiment. The subsequent analyses were performed using MATLAB.

To examine the time course of learning during the visuomotor associative learning task, we calculated the mean correct rate and 95% confidence interval for every 10 trials under both the learning and over-trained conditions (Figure 1C). For the uncertain condition, we calculated the mean correct rate and 95% confidence interval for every 10 trials when the learned and random objects were presented (Figure 1F). Next, to examine the behavioral strategies of the monkeys, we analyzed their differences in behavior between trials, after the correct and erroneous trials, when the same objects were presented. Specifically, for trials in which the same object was presented, the probability of same-direction saccades in trials following erroneous ones was subtracted from that of subsequent correct trials. We then calculated and compared these values for the first (first) and second half (second) of each session (Figures 1D and 1G; Wilcoxon rank-sum test *p* < 0.05).

To analyze the responses to visual stimuli, we performed principal component analysis (PCA) on all recorded dentate nucleus neurons (Figure S1). First, we subtracted the mean firing rate over a period of 1–300 ms before the presentation of the fractal object from the firing rate, over a period of 1–400 ms after the presentation of the fractal object. Then, we divided the 400 ms corrected neural activity by the analysis window size of 50 ms and calculated the mean over the eight analysis windows. A matrix of 1008 (number of neurons) × 8 (number of analysis windows) was prepared, for which the eigenvectors and eigenvalues were obtained via PCA, using the pca function in MATLAB.

For quantification, we measured the firing rate during the following periods: 1) 1–300 ms before fractal object presentation (baseline period), 2) 1–150 ms after fractal object presentation (early visual period), and 3) 201–400 ms after fractal object presentation (late visual period). Neurons that showed significant differences between neural activity during the baseline period and either the early or late visual periods (determined via one-way ANOVA followed by Scheffé’s method, p < 0.01) were used for subsequent analyses. The neurons with significant firing modulation during the early visual period in the learning or over-trained conditions were defined as phasic-response neurons (Figure 3E), whereas those that fired during the late visual period were defined as sustained-response neurons (Figure 3F). For both types, activity during the early or late visual period (respectively) was measured and used in subsequent quantitative analyses. The time course of neuronal activity for each condition was qualitatively examined by computing the spike density function using a Gaussian kernel (σ = 15 ms).

To investigate the direction selectivity of the sustained-response neurons, we compared the visual response during the late visual period to contralateral and ipsilateral saccade objects in the correct trials (Figure 4).

Neurons that exhibited prominent visual responses to the ipsilateral and contralateral saccade objects during the late visual period were defined as “Ipsilateral-object preferred neurons” (Figures 4A and 4C) and “Contralateral-object preferred neurons” (Figures 4B and 4D), respectively. Direction selectivity was calculated by dividing the activity to the ipsilateral saccade object, minus that to the contralateral saccade object by the sum of the activity to the ipsilateral saccade object and that to the contralateral saccade object (Figure 4E).

## Data Availability

The behavioral and neural data obtained in this study are available from the corresponding author upon reasonable request.

## Code Availability

The code used to analyze the data collected in this study is available from the corresponding author upon reasonable request.

## Acknowledgments

We would like to thank S. Nishino and N. Kajiwara for performing the animal care for this study, Y. Suwa and A. Yokoyama for their administrative support, K. Kobayashi for manufacturing the equipment, and all of the lab members for their comments and discussions. The Japanese monkeys were provided by the National Bio-Resource Project. This work was supported in part by grants from JST, PRESTO (grant numbers: JPMJPR21S4; MEXT, 21H05800 and 21H05036), and the Takeda Science Foundation.

## Author Contributions

Y.A. and J.K. designed the study, performed the experiments, and analyzed the data.

Y.A. and J.K. wrote the manuscript, with contributions from H.Y. and M.M.

## Additional information

Competing financial interests: The authors declare no competing financial interests related to this work.

## Notes

### Competing Interest Statement

The authors have declared no competing interest.

