## Supplemental Figure 1 for "Sustained visual signals in the primate cerebellar dentate nucleus drive associative learning"

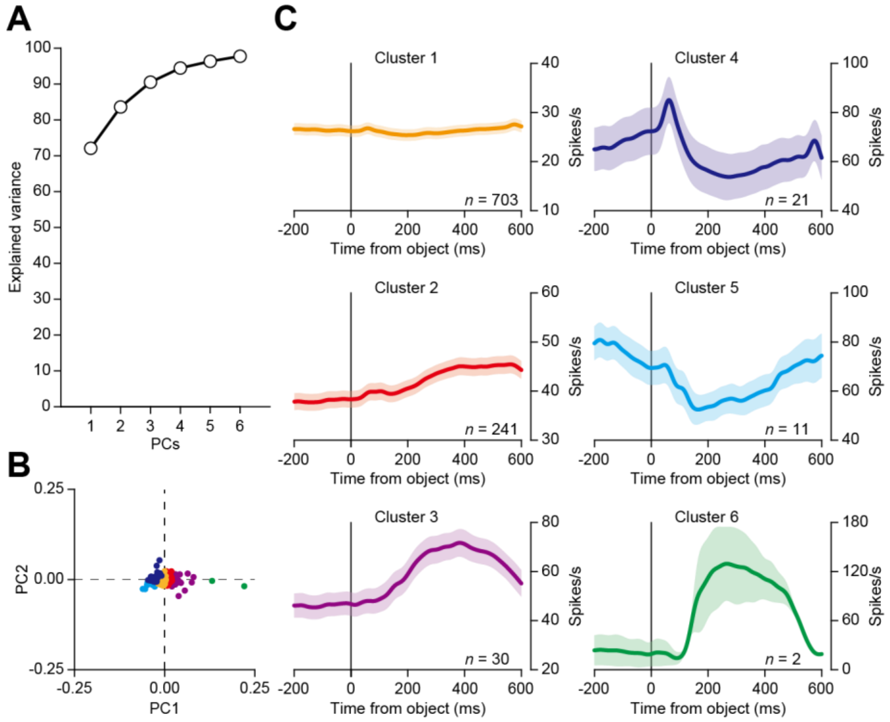


**Figure S1. Principal component analysis of dentate nucleus neurons.**

(A) Cumulative variance explained via PCA in the responses of dentate nucleus neurons to visual stimuli under the learning condition. PC1: 72.1%, PC2: 11.5%, PC3: 6.9%, PC4: 3.9%, PC5: 1.9%, and PC6: 1.4%. (B) Principal component scores of PC1 and PC2. Each point indicates the principal component score of one neuron, and the color of each point indicates the six clusters, defined as PC1–PC6 (PC1: orange, PC2: red, PC3: purple, PC4: dark blue, PC5: light blue, PC6: green). (C) Time courses of the population-level neuronal activity under the learning condition. The color of each trace corresponds to the colors of the points in Figure S1B, and indicates the population-level activity of the neurons in each cluster.
